# On the distribution of tract lengths during adaptive introgression

**DOI:** 10.1101/724815

**Authors:** Vladimir Shchur, Jesper Svedberg, Paloma Medina, Russ Corbett-Detig, RASMUS Nielsen

## Abstract

Admixture is increasingly being recognized as an important factor in evolutionary genetics. The distribution of genomic admixture tracts, and the resulting effects on admixture linkage disequilibrium, can be used to date the timing of admixture between species or populations. However, the theory used for such prediction assumes selective neutrality despite the fact that many famous examples of admixture involve natural selection acting for or against admixture. In this paper, we investigate the effects of positive selection on the distribution of tract lengths. We develop a theoretical framework that relies on approximating the trajectory of the selected allele using a logistic function. By numerically calculating the expected allele trajectory, we also show that the approach can be extended to cases where the logistic approximation is poor due to the effects of genetic drift. Using simulations, we show that the model is highly accurate under most scenarios. We use the model to show that positive selection on average will tend to increase the admixture tract length. However, perhaps counter-intuitively, conditional on the allele frequency at the time of sampling, positive selection will actually produce shorter expected tract lengths. We discuss the consequences of our results in interpreting the timing of the introgression of EPAS1 from Denisovans into the ancestors of Tibetans.

## 1. INTRODUCTION

Admixture—a process wherein genetically distinct populations hybridize and exchange alleles—is increasingly recognized as a dominant force in evolutionary genomics. In particular, admixture has the potential to introduce adaptive alleles from a donor population to a recipient, particularly if the donor population has already adapted to *e.g.*, unique environmental conditions. For example, when the ancestors of modern Ethnic Tibetans first colonized the extremely high elevation Tibetan Plateau their success in adapting to these harsh conditions stemmed in part from genes acquired from a Denisovan-like ancestral population ([12]). More generally, as genome sequence data continues to accumulate, it is increasingly clear that introgression has played a major role in shaping adaptive outcomes across a wide range of species (*e.g.* [13, 27]). However, the specific genetic signature of adaptive introgression and its impact on patterns of genetic variation remains less explored (but see *e.g.*, [33]). There is, therefore, a clear need to develop a theoretical framework to investigate the impacts of adaptive introgression on the genomes of natural populations.

When two ancestral populations hybridize, the genome of each individuals is a mixture of the ancestral populations, and the ancestry at each site in the genome can be traced back to a single population. Because alleles from a given ancestral population are initially on the same chromosomes, and therefore strongly linked within the admixed population, this process can dramatically shape linkage disequilibrium (LD). LD, and the haplotypic analog, ancestry tracts, which are unbroken stretches of sites from one ancestral population, have become the key elements of genomic inference in studying admixed populations. These two features of genetic variation are the primary units of demographic model inference within admixed populations. For example, the decay of LD [25] and the ancestry tract length distribution [7], [2] are commonly used for timing the onset of admixture assuming a neutral model during admixture. For example, in [34], alleles private to Papuans relative to European and African populations were used to call the Denisovan-like and Neanderthal-like ancestry tracts in admixed Papuan genomes.

[10], [23], [24] showed recombination, genetic drift, and migration processes act as linear operators on the 2-loci and 3-loci linkage disequilibrium (as defined in [19] and [35] respectively). In contrast, natural selection is not a linear operator on LD, and therefore approaches that assume a neutral admixture model cannot be extended to accommodate the impacts of natural selection. Therefore, another method is required to analyse the impact of selection, in particular in cases of adaptive introgression. A recent study [30] derives the ancestry tract length distribution under an infinitesimal selection model. Such a model assumes that there are many different sites and each of them is under weak selection. However, another potent force that affects admixed populations is adaptive introgression, where one or a few adaptive genes is acquired via hybridization and each has potentially strong effects on fitness. The analysis of adaptive introgression and its impacts on linkage disequilibrium and the ancestry tract length distribution is therefore a largely unexplored topic and requires a novel theoretical framework.

The distribution of tract lengths during adaptive introgression is closely related to the study of the strength of linkage disequilibrium. Two main models that address this question are 2-loci and 3-loci linkage models, which model the joint behaviour of 2 and 3 linked loci respectively. Substantial theory and numerical simulations are known from earlier works (e.g. [18], [37], [38], [19], [4], [14], [5], [15], [16], [20], [21], [6]). There are two main differences between these works and our approach. The first is the problem itself: in the aforementioned works the object of interest is primarily the strength of LD in different selection scenarios, so all the sites are considered to be segregating in a single ancestral population, and allele frequencies at all the sites affect the strength of LD. In our work we are instead interested in the population ancestry of a locus in a proximity of a selected site. The second difference is that these works consider forward time models, and we work under backward time coalescent.

The distribution of ancestry tract lengths around a locus under selection can reveal the strength of selection and the timing of admixture without requiring the assumption of neutrality. In this work, we present an efficient way to calculate the expected ancestry tract length distribution, and we analyse the dependence of this distribution on a range of different biological parameters. In particular, it is evident that for a given admixture proportion, selection results in longer tract lengths of introgressed segments. On the other hand, when conditioning on the allele frequency at the time of sampling, positive selection actually results in shorter ancestry tracts during adaptive introgression.

## 2. THEORY AND METHODS

### 2.1. Conceptual Overview

We model the tract length distribution around a selected locus. In other words, we want to find the probability that an ancestry tract ends at a certain distance from the selected locus because of an ancestral recombination event within the admixed population. We imagine a stochastic process along the length of the chromosome where transitions between ancestries at a given locus are caused by ancestral recombination (at this locus) such that ancestral chromosomes on the left and on the right of the recombination site are derived from different ancestral populations. As shown in [38], the strength of linkage disequilibrium between two neutral loci depends on the distance of the region containing those loci from the selected site. Similarly, transition rates between ancestry types are not uniform in recombination units with regards to the distance from the selected site.

Indeed, if a neutral locus is far away from the selected locus, the surrounding genomic region is nearly independent of the impact of selection, and the transitions between the ancestry types should occur almost as expected under neutrality. On the other hand, if a given site is very close to the selected locus, it would be strongly linked to it, and the probability of transitions to a different ancestry would be affected by the allele frequency changes in the selected locus. Because a 2-locus framework cannot model this dependency, it does not provide enough information regarding the local transition rates in genomic regions proximal to a selected site. Instead, in this work, we use a 3-locus model to calculate transition rates between different ancestries in two loci as a function of the distance to a third selected locus.

### 2.2. Selected allele trajectory

Let *A/a* denote a site under selection with *A* standing for the selected allele. We are interested in the scenario when allele *A* is introduced into the population through an adaptive introgression event that includes instantaneous replacement of a given proportion of individuals, the admixture fraction, in the recipient population. Such an introgression event is common termed an “ancestry pulse” in related works (*e.g.* [7], [2]). We assume that allele *A* was fixed in the donor population, and that prior to admixture, allele *a* was fixed in the recipient population.

The expected trajectory of an allele under selection can be well approximated using a logistic function under the following conditions. First, selection should be strong enough (*N*_*e*_*s >>* 1, where *N*_*e*_ is effective population size, and *s* is selection coefficient such that an individual with two selected alleles has fitness 1 + *s* and a heterozygous individual has fitness 1 + *s/*2). Second, the frequency of the selected allele is above a critical threshold, so in our model, the admixture fraction must not be too small. Finally, the time since adaptive introgression is not too large compared to the selection coefficient, so that the frequency of *A* is not too close to 1 at the time of sampling [36], [28].

Approximating the allele frequency trajectory of the adaptive allele using a deterministic logistic trajectory allows us to avoid integration over the stochasticity caused by genetic drift, which makes our approach similar to the mean field theory widely used in physics ([40]) and epidemiology ([17]). We show that this approximation is highly accurate, in a certain range of parameters, for estimating the expected tract lengths. In this range, the expected tract length can be considered invariant under the order of integration over genetic drift and over the tract length distribution.

Outside this range, when the proportion of introgression is very small, which means that the drift is strong relative to selection, the logistic function will yield inaccurate predictions. In this case the expected trajectory can be estimated by generating a large random sample of stochastic trajectories. This can be done efficiently, because in practice the selection coefficient *s <<* 1 is small, and the diploid population can be approximated by a haploid population under genic selection. At each generation the selected allele frequency is simulated from a binomial distribution with probability weighted by the total fitness contributed of the selected allele. We average allele frequencies at each generation over the simulated trajectories (conditioning on observing the selected allele at the time of sampling). We will show that such an approach vastly extends the applicability of our method to cases that require greater stochasticity. In particular, we used it to analyse the Denisovan introgression into ethnic Tibetans, where the proportion of introgression is estimated to be as small as 0.06% [12], and therefore is outside the range of parameters where the logistic approximation to the allele frequency trajectory yields accurate results.

### 2.3. 3-locus model during adaptive introgression

Let *b* and *c* be two neutral loci near *A/a* so that we have a 3-loci segment *A/a − b − c*, where loci *A/a, b* and *c* follow each other sequentially on the chromosome. Let *r*_1_ be the distance (in Morgans) between *A/a* and *b, r*_2_ is the distance between *b* and *c*. We want to know the ancestry of loci *b* and *c*. In other words, we want to track the lineage leading to each locus to the time of introgression, while simultaneously tracking how recombination and coalescence act on the segment. If at the time of introgression the locus is on a haplotype carrying *A*, then, by definition, its ancestry is from the introgressed population (denoted ancestry type 1). If, in contrast, the locus is linked to the allele *a*, then it comes from the recipient population (denoted ancestry type 0). Ancestry type 0 corresponds to the recipient population and ancestry type 1 corresponds to the donor population.

The importance of this 3-locus model is that we can calculate transition rates between ancestry types, and hence in particular, we can numerically calculate the distribution of tract lengths of ancestry type 1 (or type 0) near the site *A/a*.

### 2.4. Model derivation

Under the coalescent with recombination, the model can be considered Markovian backwards in time, when conditioned on the allele frequency path. To describe the dynamics of the adaptive introgression 3-locus model, we need to enumerate the possible states, which describes the ancestry configurations across the three loci, and we need to find the transition rates between them.

The model has 6 possible states, each representing an ancestral configuration for a chromosome with 3 sites: *A/a, b* and *c*. We use an asterisk to indicate alleles in sites *b* and *c* which are not ancestral on the chromosome of interest.

- (*A −b − c*): both loci *b* and *c* are on the same chromosome carrying allele *A*,
- (*A −b − *, A− *− c*): loci *b* and *c* are on different chromosomes, both carrying allele *A*,
- (*A −b−*, a −*− c*): loci *b* and *c* are on different chromosomes, and the chromosome with locus *b* is carrying allele *A*, while the chromosome with locus *c* is carrying allele *a*,
- (*a −b −*, A−*− c*): loci *b* and *c* are on different chromosomes, and the chromosome with locus *b* is carrying allele *a*, while the chromosome with locus *c* is carrying allele *A*,
- (*a − b− *, a− *− c*): loci *b* and *c* are on different chromosomes, both carrying allele *a*,
- (*a − b − c*): both loci *b* and *c* are on the same chromosome carrying allele *a*.

We denote the frequency of the selected allele at time *t* by *ω*(*t*). As we indicated previously, we assume that *ω*(*t*) deterministically follows a logistic function:

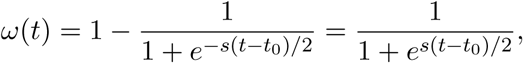

because we are working in backward time. This is a reflection of a logistic function relative to *t* = 0. The time of introgression corresponds to a point, *t*_1_, on the deterministic allele frequency trajectory such that *ω*(*t*_1_) equals the admixture proportion *ω*_1_ (see Figure 1). Notice, that the time of sampling *t*_0_ = *t*_1_ − *T*, where *T* is the time since introgression, does not necessarily equal *t* = 0.

**FIGURE 1.**
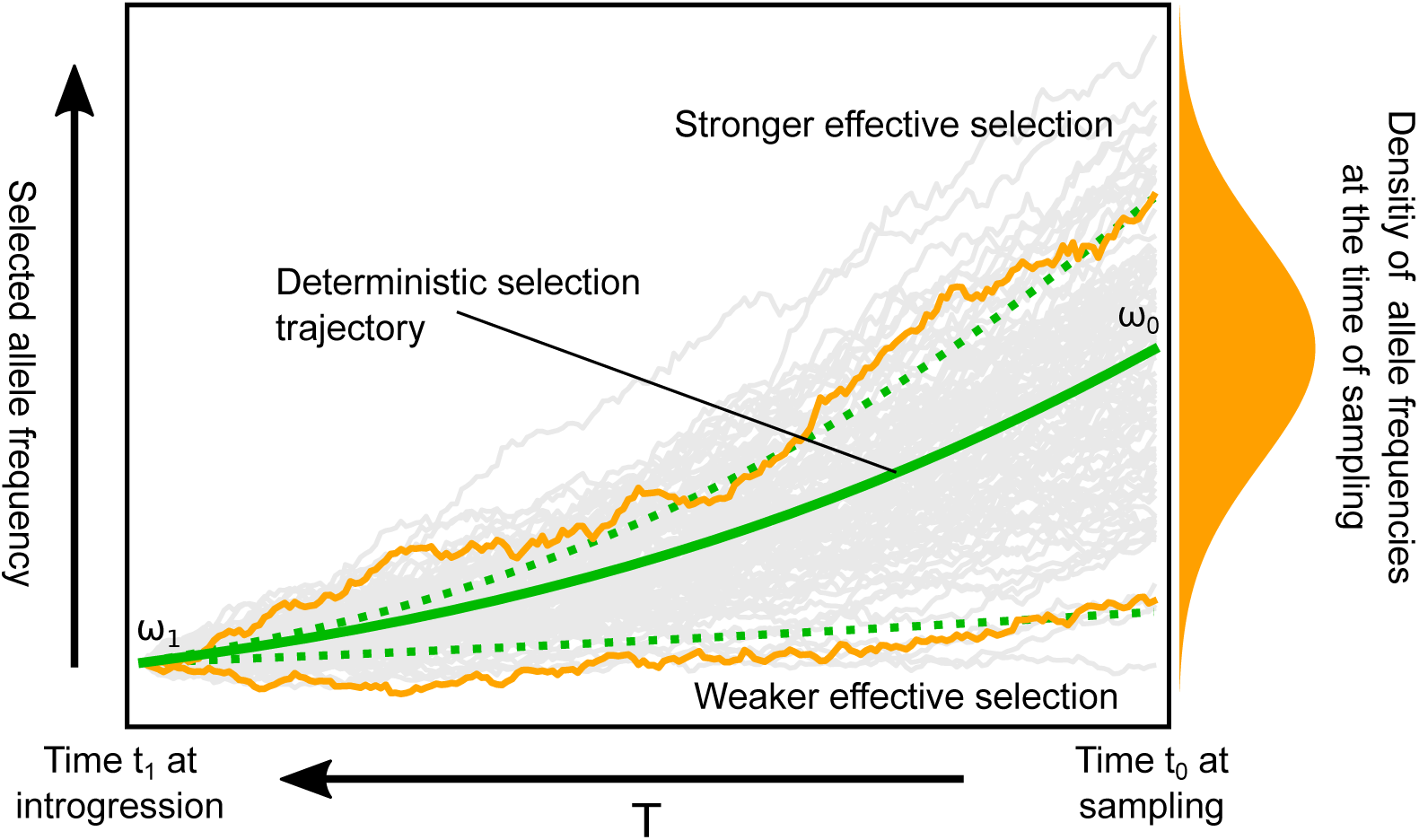
The solid green line is the expected trajectory, which is used in our deterministic model described by Equation 1. Gray lines are stochastic trajectories, which induce the distribution of observed allele frequency (orange on the right-hand side), and this distribution can be estimated from simulations. The two orange stochastic trajectories correspond to paths with stronger and weaker effective selection, and the two dotted green lines are logistic trajectories corresponding to those values of effective selection coefficients.

Recombination acts at a rate proportional to the recombination distances between loci. If a haplotype has the ancestry configuration state *A − b − c* at time *t*, then recombination between *A* and *b* occurs at rate *r*_1_ and recombination occurs between *b* and *c* at rate *r*_2_. In other words, we measure distance between the loci in Morgans. If recombination occurs between *A* and *b*, then loci *b* and *c* remain on the same chromosome. The allele at the site *A/a* on this chromosome will be *A* with probability *ω*(*t*) (state does not change) and *a* with the probability 1 *− ω*(*t*). Hence, the transition rate from state (*A − b − c*) to the state (*a −b − c*) is *r*_1_(1 *− ω*(*t*)) at time *t*. If a recombination event occurs on a chromosome with haplotype *A −b − c* between loci *b* and *c*, then it is split into two chromosomes, and there are two different ancestry configurations. In both configurations the first ancestral chromosome has haplotype *A −b − \**. The second chromosome is either *A − * −c* with probability *ω*(*t*), or *a − * −c* with probability 1 − *ω*(*t*). The other type of transition in our model is coalescence, which is possible between chromosomes that carry the same allele at the selected locus at the rate *λ* = 1*/*2*N*_*e*_, where 2*N*_*e*_ is the haploid effective population size. The full transition matrix is

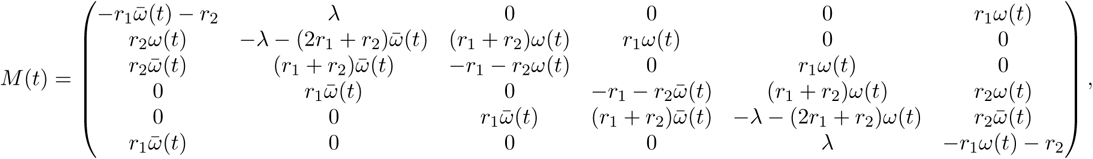

where 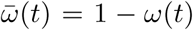 is the allele frequency of allele *a*, and the equation describing the probability of being at a certain state at time *t* is

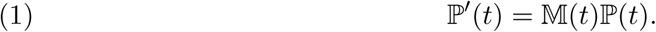

The initial condition for this differential equation is

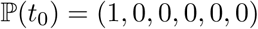

for the dynamics of introgressed tracts (hence, carrying allele *A*), and

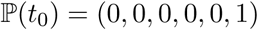

for the tracts from the recipient population (individuals with allele *a*).

Equation 1 cannot be solved analytically, because eigenvalues of the matrix *M* (*t*) cannot be derived analytically in general. However, it is easy to solve it with standard numerical methods.

### 2.5. Transition rates between ancestry of type 0 and type 1

As previously discussed, we are interested in transitions of ancestry in loci near the introgressed selected allele. More precisely, we seek the distribution of lengths of ancestry tracts that contain *A*, i.e., the set of ancestry tracts with ancestry type 1 at the selected site. In the given model the probability that a locus is of ancestry type 1 or type 0 is equal to the probability that the ancestral chromosome carries the allele *A* or *a* respectively at time of introgression *t*_1_. In the previous subsection we described a Markovian process (with regards to the backward time) under which we can calculate those probabilities.

Now we consider a new Markov process, which describes the ancestry at a locus while moving along the chromosome away from the selected locus. The states of this Markov process are again ancestries of type 0 or 1. By definition, transition rate between states *s*_1_ and *s*_2_ at position *r* of a Markov process *S*(*r*) is

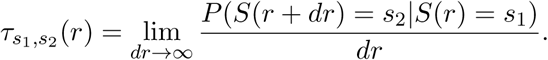

The transition rate is *τ*_10_(*r*) between ancestries of type 1 and type 0, which corresponds to recombination breaking the introgressed ancestry tract, is

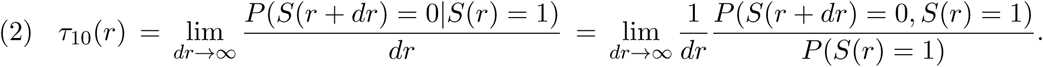

The numerator *P* (*S*(*r* + *dr*) = 0, *S*(*r*) = 1) is the probability of being at the third state (*A − b− \**, *a− * − c*), and the denominator *P* (*S*(*r*) = 1) is the marginal probability of ancestry type 1, hence it is the sum of probabilities of the first three ancestral configurations (*A− b − c*), (*A− b − *, A− * − c*) and (*A− b − *, a − * − c*). Expression 2 is easy to evaluate numerically by considering sufficiently small values of *r*_2_.

Calculation of the expected tract length for 100 introgression scenarios takes just 106 seconds for a Python implementation executed on MacBook Pro (2.9 GHz Intel Core i5) when using the logistic function to approximate the allele frequency trajectory. Transition rates are evaluated at 1000 points (different values of *r*). The code is available at the GitHub repository associated with this project https://github.com/vlshchur/DAIM.

## 3. RESULTS

### 3.1. Accuracy of approximation

In order to estimate how accurate our approximation is, we compared our results with the average tract length estimated from simulations using the forward-in-time simulation framework SELAM [1]. Briefly, we used the software to simulate a Wright-Fisher population of constant size 5000 hermaphroditic diploid individuals. We simulated a single chromosome of length one Morgan with a single selected site at position 0.5 Morgans. At each sampling point, we extracted 50 individuals (100 chromosomes) at random from within the population and output the ancestry across each chromosome. We performed 10 to 100 thousand replicates of each combination of selective coefficient, times since admixture and admixture proportion.

We simulated both relatively weak and strong selection, and we allowed selection to act for different periods of time. In all of the scenarios that we explored, the relative error does not exceed 5%, and in more than half of all cases it is within 2% (Tables 1 and 2). This level of precision should be sufficient for most applications given that the uncertainty in the real data analysis is usually much greater than the error of the approximation model.

**TABLE 1.** The accuracy of the deterministic approximation for the expected tract length under adaptive introgression compared to estimates from simulations. For every set of parameters (introgression fraction, selection coefficient and time of introgression), we performed 100 thousand replicates of simulations in each case. The haploid effective population size was 10000 chromosomes, with 100 chromosomes sampled from each population. The relative error was calculated by comparing the simulated expected tract length to the prediction given by the deterministic model.

| Introgression parameters |  |  | Expected tract length |  |  | Relative error |
| --- | --- | --- | --- | --- | --- | --- |
| Proportion | Selection | Time | Simulations | Deterministic approximation | Weighted deterministic approximation |  |
| 0.01 | 0.001 | 50 | 0.0417436 | 0.0401907 | 0.04027677 | 3.7% |
|  |  | 100 | 0.0197123 | 0.0200985 | 0.02014723 | 2.0% |
|  |  | 500 | 0.00407317 | 0.00402525 | 0.004044527 | 1.1% |
|  |  | 1000 | 0.00206252 | 0.00201644 | 0.0020327 | 2.2% |
| 0.01 | 0.01 | 50 | 0.0415686 | 0.0402287 | 0.04029757 | 3.3% |
|  |  | 100 | 0.0202129 | 0.02014 | 0.02017217 | 0.4% |
|  |  | 500 | 0.00423004 | 0.00411598 | 0.004121694 | 2.8% |
|  |  | 1000 | 0.00237283 | 0.00227902 | 0.002258553 | 4.0% |
| 0.05 | 0.001 | 50 | 0.0427474 | 0.0419025 | 0.04187598 | 2.0% |
|  |  | 100 | 0.0210069 | 0.0209646 | 0.020949 | 0.2% |
|  |  | 500 | 0.00428813 | 0.00421551 | 0.004214419 | 1.7% |
|  |  | 1000 | 0.00217073 | 0.00212356 | 0.00213722 | 2.2% |
| 0.05 | 0.01 | 50 | 0.0428456 | 0.0420981 | 0.04206946 | 1.7% |
|  |  | 100 | 0.021438 | 0.0211766 | 0.02115895 | 1.2% |
|  |  | 500 | 0.00470731 | 0.00462923 | 0.004616305 | 1.7% |
|  |  | 1000 | 0.00290033 | 0.00286404 | 0.002865537 | 1.3% |

**TABLE 2.** The accuracy of the deterministic approximation for the expected tract length under adaptive introgression compared to the estimates from simulations. * For these scenarios, which include relatively small admixture fractions, the logistic function does not accurately describe the allele frequency trajectory, so we numerically estimated the mean trajectory using stochastic simulations.

| Introgression parameters |  |  | Expected tract length |  |  |
| --- | --- | --- | --- | --- | --- |
| Proportion | Selection | Time | Simulations | Deterministic approximation* | Relative error |
| 0.0006 | 0.01 | 1500 | 0.00176 | 0.00173 | 1.7% |
|  |  | 1750 | 0.00158 | 0.00160 | 1.3% |
|  |  | 2000 | 0.001503 | 0.001500 | 0.2% |
|  |  | 2250 | 0.00141 | 0.00142 | 0.7% |
| 0.0006 | 0.02 | 1500 | 0.00234 | 0.00230 | 1.7% |
|  |  | 1750 | 0.00222 | 0.00215 | 3.2% |
|  |  | 2000 | 0.00210 | 0.00203 | 3.3% |
|  |  | 2250 | 0.00205 | 0.00193 | 5.9% |
| 0.025 | 0.001 | 1500 | 0.001415 | 0.001436 | 1.5% |
|  |  | 1750 | 0.001227 | 0.001244 | 1.4% |
|  |  | 2000 | 0.001136 | 0.001101 | 3.1% |
|  |  | 2250 | 0.001020 | 0.000990 | 2.9% |
|  |  | 2500 | 0.000883 | 0.000902 | 2.2% |
| 0.025 | 0.005 | 1500 | 0.00164 | 0.00158 | 4.9% |
|  |  | 1750 | 0.00146 | 0.00142 | 2.7% |
|  |  | 2000 | 0.00133 | 0.00130 | 2.3% |
|  |  | 2250 | 0.00126 | 0.00121 | 4.0% |
|  |  | 2500 | 0.00118 | 0.00114 | 3.4% |

In our approach the allele frequency at the time of sampling *ω*_0_ is uniquely defined by the parameters of introgression (selection coefficient *s*, time since introgression *T* and the admixture fraction *ω*_1_). In reality, due to the effect of genetic drift, *ω*_0_ is a random variable (see Figure 1).

In order to take into account the uncertainty of *ω*_0_, we developed the following approach: first, for a given scenario of introgression we empirically estimated the distribution, 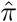, of allele frequency at the time of sampling. To do so, for each set of parameters *s, T* and *ω*_1_, we simulated 100,000 replicates to obtain a distribution of *ω*_0_. For each simulation, we obtained a sample of 100 chromosomes, so possible values of the allele frequencies at the time of sampling are 0.01, 0.02, *…*, 0.99 (we exclude cases with no variation in the sampled chromosomes, that is with sample allele frequencies of 0 and 1). Second, for each value, 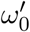, of the allele frequency at the time of sampling, we found an effective selection coefficient 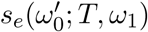, such that the expected allele frequency at the time of sampling (given by the logistic trajectory) for parameters *s*_*e*_, *T* and *ω*_1_ is 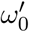. The effective selection coefficient is given by the following equation

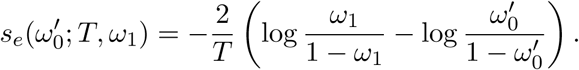

For each such set of parameters we calculated the expected tract length 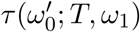 as the expectation of *τ* integrated over simulated values of *ω*_0_. As shown in table 1 (column “weighted deterministic approximation”), the resulting estimates of expected tract lengths are still consistent with simulations and the deterministic approximation.

As previously discussed, the logistic function is a good approximation for the allele frequency trajectory only when selection is strong. Still, we wondered if the deterministic approach would give consistent results for a case of neutral admixture without adaptive introgression. Following Liang, Nielsen [22] the expected tract length under a neutral admixture model can be calculated as

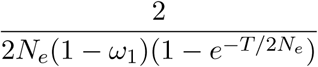

under the SMC’ model [26]. Notice that there is a factor of two in the numerator due to the fact that we condition on observing the introgressed allele on a haplotype. We find that in the limiting case of no selection, the approximation closely approximates the analytical result from [22] (Table 3).

**TABLE 3.**
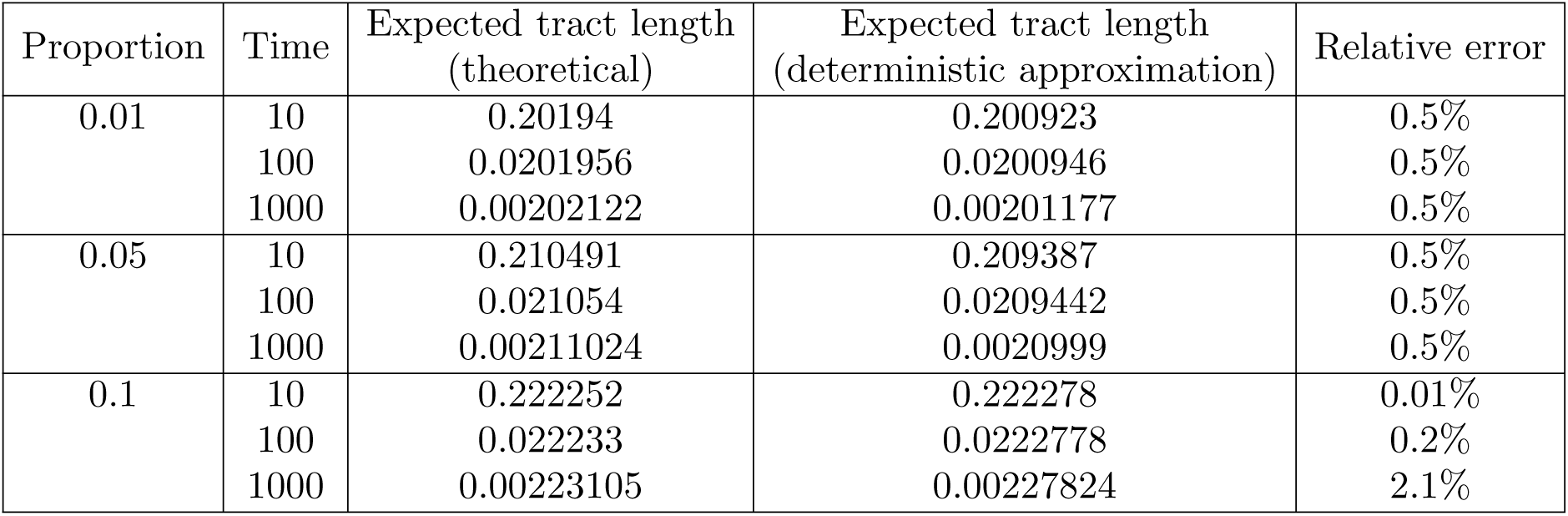
Our deterministic prediction for the expected tract length in a neutral model is consistent with the theoretical expectation under SMC’ model.

| Proportion | Time | Expected tract length<br>(theoretical) | Expected tract length<br>(deterministic approximation) | Relative error |
| --- | --- | --- | --- | --- |
| 0.01 | 10 | 0.20194 | 0.200923 | 0.5% |
|  | 100 | 0.0201956 | 0.0200946 | 0.5% |
|  | 1000 | 0.00202122 | 0.00201177 | 0.5% |
| 0.05 | 10 | 0.210491 | 0.209387 | 0.5% |
|  | 100 | 0.021054 | 0.0209442 | 0.5% |
|  | 1000 | 0.00211024 | 0.0020999 | 0.5% |
| 0.1 | 10 | 0.222252 | 0.222278 | 0.01% |
|  | 100 | 0.022233 | 0.0222778 | 0.2% |
|  | 1000 | 0.00223105 | 0.00227824 | 2.1% |

Another case where the logistic function is not expected to be a good approximation of the mean trajectory, is when the admixture proportion is close to zero. To explore this issue, we chose a set of parameters, which are similar to Neanderthal introgression into non-Africans (2.5%), and to the Denisovan introgression into Tibetans (0.06%). We found that the logistic approximation performs poorly for the Denisovan-like scenario, and is not very accurate (up to 10% relative error) for the Neanderthal-like introgression with week selection (*s* = 0.001); in particular, the predicted allele frequency is strongly underestimated. Presumably these discrepancies reflect the increased impacts of genetic drift for small admixture proportions. To address this issue, we estimated the expected trajectory by forward-in-time stochastic simulations of a large sample of possible allele frequency trajectories (see subsection 2.2 for details). The stochastically estimated expected trajectory is then used instead of the logistic function in Equation 1 to model the tract length distribution. The deterministic model (with numerically pre-computed expected trajectories) gives tract length estimates quite similar to those observed in the simulated data sets (Table 2).

### 3.2. Variance

The variance in the tract length distribution predicted by our method is an accurate approximation of the across population variance, that is, the variance among all the tracts obtained in many replicates with the same parameters (see Table 4). This is equivalent to the variance in tract length among tracts within a population if the tracts introgressed into the population independently of each other and have not coalesced or recombined with each other. Of course, in reality, two tracts might coalesce before the time of introgression, so their lengths are not independent of each other. Hence, we expect that the within-population variance is smaller than across-population variance. On the other hand, if introgression is recent enough, then the chance of coalescence is small, and the correlation between tract lengths within population will be small.

**TABLE 4.** Simulated across and within population variance and the across population variance estimated from the deterministic model. For smaller times since introgression, between and within populations variances are similar. However, for older introgression times (in the second part of the table) this is not true, and the within population variance becomes smaller as a result of the correlation induced by coalescences. The introgression with 0.0006 (or, 0.06%) admixture proportion was chosen to approximate Denisovan introgression into Tibetans. The introgression with 0.025 (or, 2.5%) admixture proportion was chosen to approximate Neanderthal introgression into non-Africans.

| Introgression parameters |  |  | Standard deviation of tract length |  |  |
| --- | --- | --- | --- | --- | --- |
| Proportion | Selection | Time | Simulated<br>(across populations) | Simulated<br>(within population<br>all replicates) | Deterministic<br>approximation |
| 0.01 | 0.001 | 50 | 0.0293966 | 0.02740539 | 0.0285128 |
|  |  | 100 | 0.013484 | 0.01337553 | 0.0142598 |
|  |  | 500 | 0.00279459 | 0.002334261 | 0.00285811 |
|  |  | 1000 | 0.00145372 | 0.001131087 | 0.00143309 |
| 0.01 | 0.01 | 50 | 0.0287187 | 0.02757148 | 0.0285337 |
|  |  | 100 | 0.0137616 | 0.01348083 | 0.0142825 |
|  |  | 500 | 0.0030498 | 0.002621121 | 0.00290384 |
|  |  | 1000 | 0.00154791 | 0.001438002 | 0.00154791 |
| 0.05 | 0.001 | 50 | 0.0304915 | 0.02958953 | 0.0297278 |
|  |  | 100 | 0.0149323 | 0.01464238 | 0.014875 |
|  |  | 500 | 0.00303497 | 0.002756722 | 0.00299345 |
|  |  | 1000 | 0.00155372 | 0.001336331 | 0.00150908 |
| 0.05 | 0.01 | 50 | 0.0306196 | 0.02973097 | 0.0298345 |
|  |  | 100 | 0.0149798 | 0.01483033 | 0.0149894 |
|  |  | 500 | 0.00324606 | 0.003119859 | 0.00319587 |
|  |  | 1000 | 0.0018881 | 0.001784578 | 0.00183546 |
| 0.0006 | 0.01 | 1500 | 0.001173 | 0.000962 | 0.001125 |
|  |  | 1750 | 0.000998 | 0.000889 | 0.001014 |
|  |  | 2000 | 0.000954 | 0.000811 | 0.000930 |
|  |  | 2250 | 0.000884 | 0.000751 | 0.000864 |
| 0.0006 | 0.02 | 1500 | 0.001423 | 0.001241 | 0.001350 |
|  |  | 1750 | 0.001312 | 0.001132 | 0.001230 |
|  |  | 2000 | 0.001221 | 0.001050 | 0.001136 |
|  |  | 2250 | 0.001179 | 0.000974 | 0.001061 |
| 0.025 | 0.001 | 1500 | 0.001052 | 0.000806 | 0.001015 |
|  |  | 1750 | 0.000840 | 0.000677 | 0.000878 |
|  |  | 2000 | 0.000835 | 0.000589 | 0.000776 |
|  |  | 2250 | 0.000691 | 0.000535 | 0.000696 |
|  |  | 2500 | 0.000608 | 0.000477 | 0.000633 |
| 0.025 | 0.005 | 1500 | 0.001136 | 0.000991 | 0.001082 |
|  |  | 1750 | 0.000970 | 0.000875 | 0.000956 |
|  |  | 2000 | 0.000887 | 0.000784 | 0.000863 |
|  |  | 2250 | 0.000836 | 0.000726 | 0.000793 |
|  |  | 2500 | 0.000779 | 0.000672 | 0.000737 |

Indeed, for most of the scenarios with 50 and 100 generations since the time of introgression (Table 4) the variances of the tract lengths are similar. For larger times, the across-populations and within-population variance are different by up to 20%, reflecting the increased probability of coalescence among selected haplotypes that share a recombination event within the admixed population.

### 3.3. Dependence of mean tract length on different parameters

The deterministic approach facilitates the calculation of expected tract lengths for a wide array of parameters. In Figure 2 we illustrate the dependency of mean tract length on the admixture fraction given the time of introgression and selection coefficient. As expected, a larger admixture fraction results in longer expected tract lengths. In Figure 3 we illustrate the dependency on the selection coefficient given the time of introgression and admixture proportion. Again, stronger selection results in longer expected ancestry tract lengths.

**FIGURE 2.**
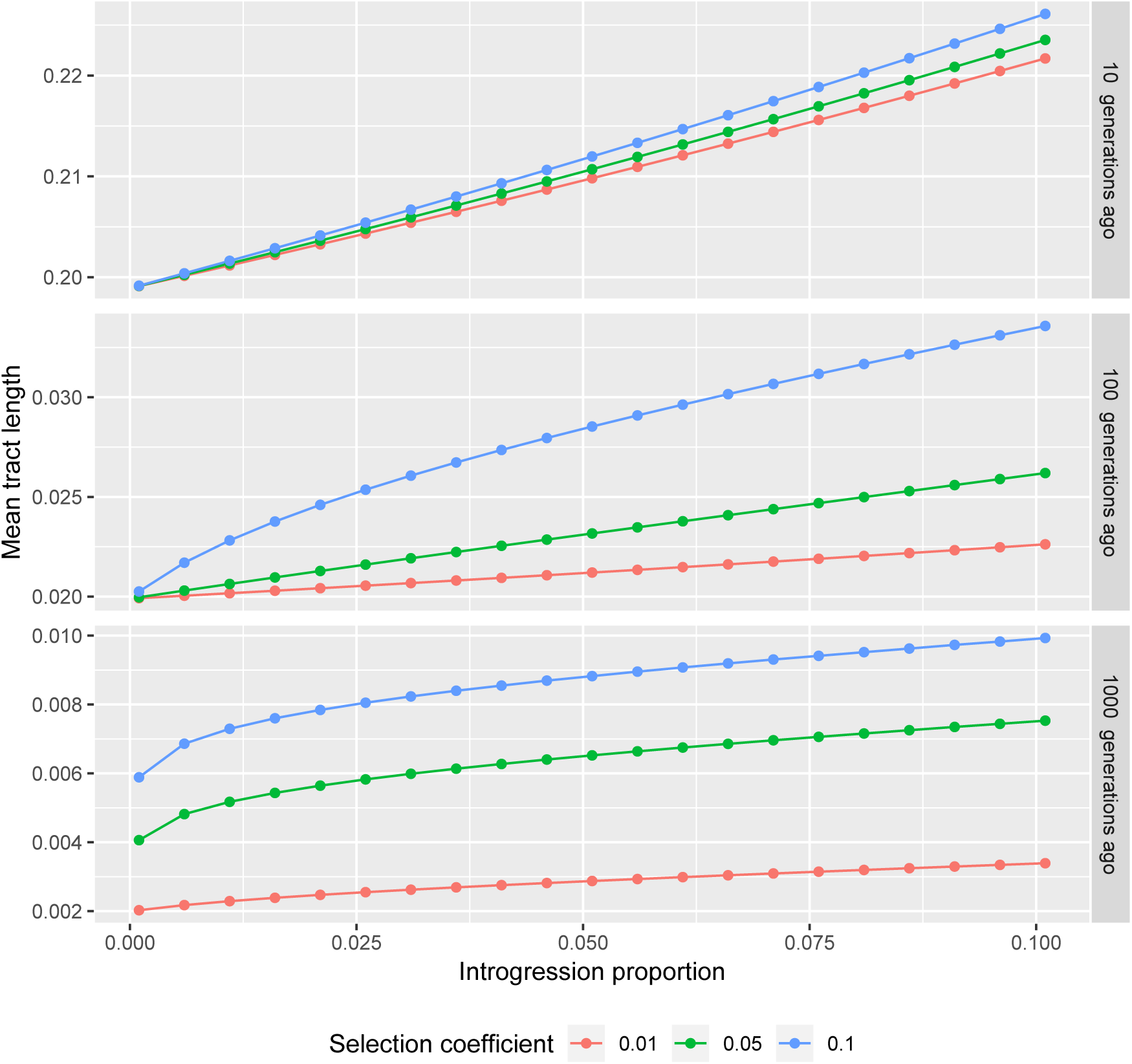
Dependence of the expected tract length on the proportion of introgression. Different panels correspond to different times of introgression (10, 100 and 1000 generations respectively). Notice that as the time since introgression changes by an order of magnitude, the tract lengths change by an order of magnitude also. So, the y-axes here are shown on different scales.

**FIGURE 3.**
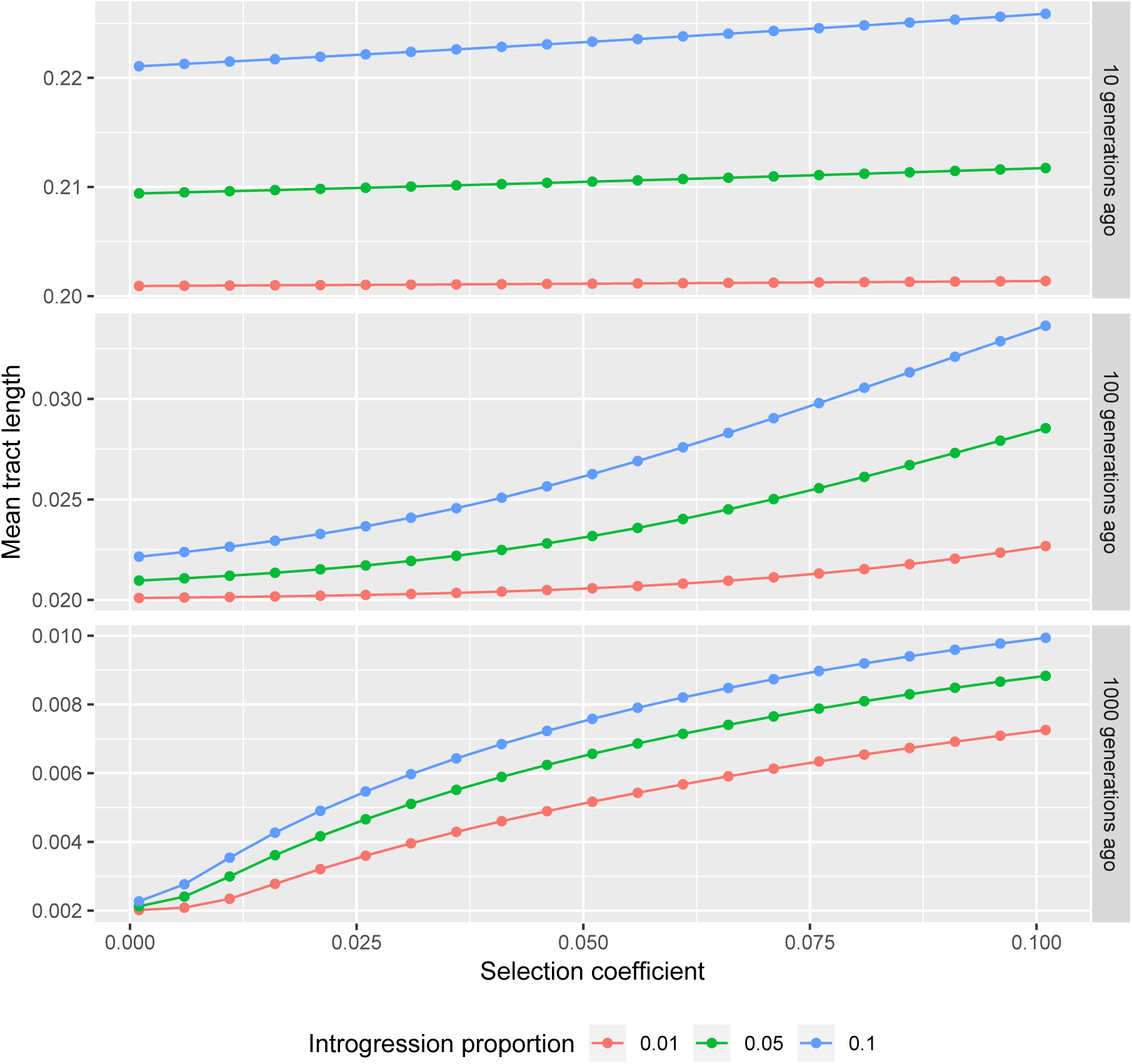
Dependence of the expected tract length on the strength of selection for different times of introgression.

The strength of selection has a stronger effect on the expected tract length than the admixture proportion. For example, consider an adaptive introgression model with time *T* = 1000, selection coefficient *s* = 0.01 and admixture proportion 0.025. The expected tract length is 0.0025 Morgans under this model. If we double the admixture proportion to 0.05, the expected tract length increases to 0.0029 Morgans, or approximately 13%). If on the other hand we increase the strength of selection by a factor of two, then the expected tract length increases by 41% and becomes 0.0036 Morgans.

The new approximation facilitates investigation of another important dependency, which would otherwise require prohibitively large numbers of simulations, namely the dependency of the expected tract length on the observed allele frequency. In Figure 4 we demonstrate that conditioning on the allele frequency at the time of sampling, stronger selection actually reduces the expected tract length. This is explained by the fact that stronger selection decreases (going backward in time) the allele frequency faster than weaker selection. Hence, the probability of recombining back to a haplotype that contains the selected allele following a recombination event is smaller for scenarios with stronger selection. More formally, assume that we compare two scenarios with selection coefficients *s*_1_ and *s*_2_ (all other parameters are the same), and that *s*_1_ *< s*_2_. Let *ω*^(*i*)^ be the logistic trajectory for selection coefficient *s*_*i*_. We condition on the observed allele frequencies at time *t*_0_, hence *ω*^(1)^(*t*_0_) = *ω*^(2)^(*t*_0_). Ancestral recombination occur on a chromosome independent of the strength of selection. Assume that recombination occurred at time *t*_*r*_ *> t*_0_. The expected allele frequency at time *t*_*r*_, which is defined by the logistic function, is larger for the smaller selection coefficient: *ω*^(1)^(*t*_*r*_) *> ω*^(2)^(*t*_*r*_). Hence, the chance that the remaining part of the chromosome coalesces back to a haplotype that contains the selected allele is higher for *s*_1_.

**FIGURE 4.**
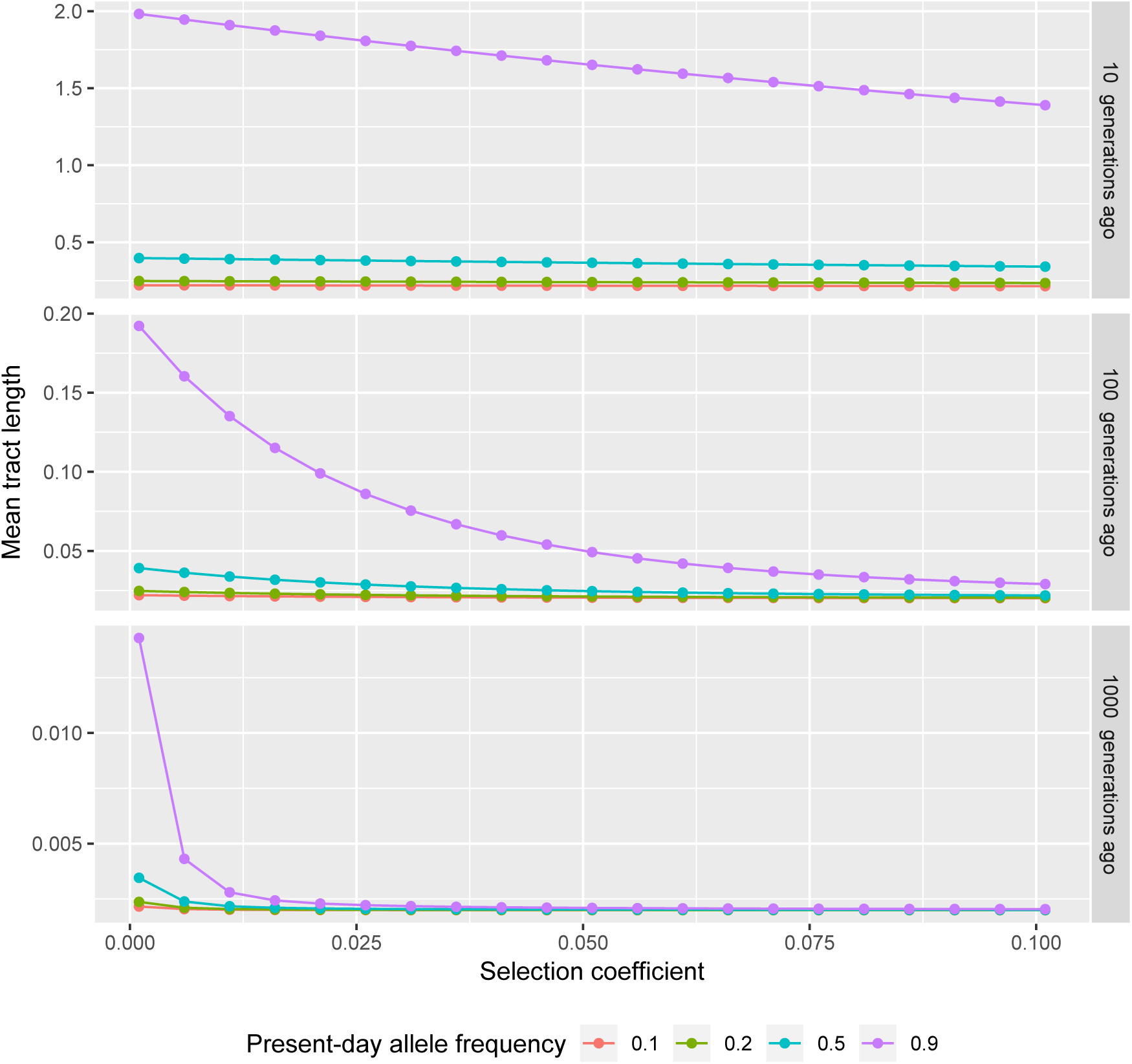
Dependence of expected tract length on the strength of selection conditioned on the allele frequency at the time of sampling.

### 3.4. Denisovan introgression into Tibetans

A well-known example of adaptive introgression is Denisovan introgression into Tibetans which facilitated altitude adaptation by introducing an adaptive allele of EPAS1 affecting red blood cell production into the ancestors of modern Tibetans [41], [12]. We wondered how selection might affect estimates of the time of introgression, conditioning on the present-day allele frequency of EPAS1 of 85% [12] and an overal genomic admixture proportion of *ω*_1_ = 0.06 ± 0.03% [32].

The admixture proportion is very small, and the logistic function will therefore be quite inaccurate. For a range of values of selection coefficient (*s* = 0.005, 0.006, *…*, 0.02), we numerically estimated the expected trajectories (using the stochastic trajectory simulator) for the given admixture proportion. Given the frequency of the adapted allele in the modern population, the time since introgression is estimated as the time needed for the allele to reach this observed allele frequency following the expected trajectory.

We summarise the dependency of the introgression time on the strength of selection in Table 5 for the mean value *ω*_1_ = 0.06%. It is clear from the table that dating the time of introgression from the tract length under the hypothesis of neutrality, would cause underestimation of the time of introgression by about 20 − 25%. For example, if the estimated tract length is 0.00117 Morgans, the corresponding introgression time without selection affecting the allele frequency is 1791 generations, or about 45000 years. However, taking selection into account, this tract length corresponds to *s* = 0.006 Morgans and an introgression time of 2282 generations, or 57000 years ago. Therefore, analyses that seek to understand the timing of introgression events of adaptively introgressed alleles will be greatly facilitated by incorporating the dependency on the strength of selection on the expected tract length distribution.

**TABLE 5.**
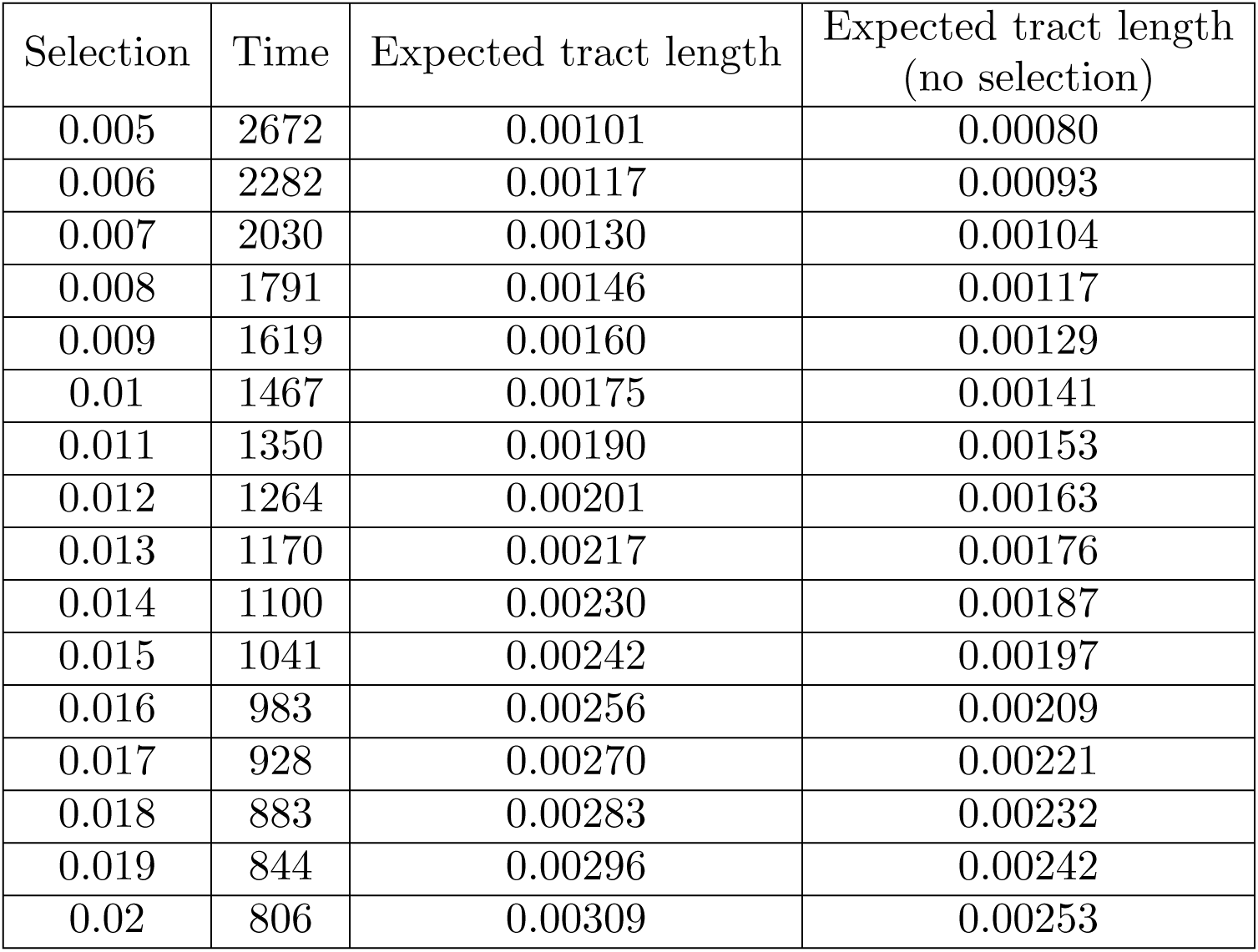
The effect of assumptions regarding the strength of selection on estimates of the time of the Denisovan introgression into Tibetans for EPAS1. We assume an initial introgression fraction of 0.06% [32], and a present-day allele frequency for the EPAS1 allele of 85% in Tibetans [12]. The value of the selection coefficient then determines the method of moments estimate of the time since introgression.

## 4. CONCLUSIONS AND PROSPECTS

Adaptive introgression is an important and common phenomenon in evolutionary genetics [9]. In this paper we derived an accurate approximating mathematical model for the ancestry tract length distribution under adaptive introgression near a selected site. This framework facilitated our efficient calculation of the distribution of tract lengths under a range of plausible adaptive introgression scenarios. We demonstrated, perhaps counter-intuitively, that conditioned on the observed allele frequency, stronger selection produces shorter admixture tracts. Furthermore, as an example, we showed that selection should be taken into account when dating the well-known case of altitude adaptation in Tibetans through the introgression of EPAS1 gene from Denisovans. When ignoring the impacts of selection, the time since introgression might be underestimated by as much as 20-25%.

Our results illustrate that selection should be carefully considered and incorporated when studying adaptive introgression events. More generally, this framework opens new possibilities for understanding the timing of admixture and the strength of adaptive introgression across a wide range of populations. Our work therefore lays foundational groundwork for the development of powerful inference frameworks that can detect adaptive introgression by searching for the characteristic ancestry tract length distribution we defined here. Particularly in light of the fact that adaptive introgression is common historically, and may be accelerated in the future by anthropogenic impacts such as climate change (*e.g.*, [29]), developing a strong theoretical basis for understanding the impacts of adaptive introgression on patterns of genetic variation is crucial for interpreting the genomic consequences of admixture.

## 5. ACKNOWLEDGMENT

This research was supported by NIH grant R01GM116044-01 (RN, VS) and a grant from the Danish DFF (RN). This work was also supported by NIH R35GM128932 (RC-D). PM was supported by NIH training grant T32 HG008345. This research was also supported by grant RFBR 19-07-00515 (VS).

